# *In vitro* reconstitution of Sgk3 activation by phosphatidylinositol-3-phosphate

**DOI:** 10.1101/2021.04.13.439688

**Authors:** Daniel Pokorny, Linda Truebestein, Kaelin D. Fleming, John E. Burke, Thomas A. Leonard

## Abstract

Serum- and glucocorticoid-regulated kinase 3 (Sgk3) is activated by the phospholipid phosphatidylinositol-3-phosphate (PI3P) downstream of growth factor signaling and by Vps34-mediated PI3P production on endosomes. Upregulation of Sgk3 activity has recently been linked to a number of human cancers. Here, we show that Sgk3 is regulated by a combination of phosphorylation and allosteric activation by PI3P. We demonstrate that PI3P binding induces large conformational changes in Sgk3 associated with its activation, and that the PI3P binding pocket of the PX domain of Sgk3 is sequestered in its inactive conformation. Finally, we reconstituted Sgk3 activation via Vps34-mediated PI3P synthesis on phosphatidylinositol liposomes *in vitro*. In addition to defining the mechanism of Sgk3 activation by PI3P, our findings open up potential therapeutic avenues in allosteric inhibitor development to target Sgk3 in cancer.

## Introduction

Cellular membranes are hotspots at which numerous signaling pathways controlling various aspects of cell physiology and pathology converge. One of the most intensively studied pathways downstream of extracellular growth factors is the phosphoinositide 3-kinase (PI3K) pathway. PI3K activation by receptor tyrosine kinases, G-protein-coupled receptors, or RAS leads to the generation of various phosphoinositide lipids, that serve as a platform for the recruitment and activation of effector molecules (Martini *et al*, 2014). Dysregulation of PI3K signaling is frequently observed in cancer, diabetes, neurodegenerative and cardiac disorders, and other diseases (Fruman & Rommel, 2014; Fruman *et al*, 2017), making the pathway a prominent target for pharmacological inhibition (Marone *et al*, 2008).

Many important downstream effectors of PI3K are protein kinases. Protein phosphorylation is a ubiquitous post-translational modification that regulates cell signaling and is carried out by the arsenal of more than 500 protein kinases in the human proteome (Manning, 2002). Since the mechanism of phosphotransfer catalyzed by protein kinases is highly conserved (Endicott *et al*, 2012), individual kinases have evolved unique regulatory mechanisms to ensure that they can faithfully respond to different, often very transient, lipid-based signaling cues (Leonard & Hurley, 2011).

Serum- and glucocorticoid-regulated kinase 3 (Sgk3) [also known as cytokine-independent survival kinase (CISK)] is a serine/threonine protein kinase that is activated downstream of growth factors (Liu *et al*, 2000). Like the closely related kinase Akt, it belongs to the AGC kinase family and is activated by phosphorylation of a hydrophobic motif in its C-terminal tail by mTORC2 (Bago *et al*, 2016) and subsequent PDK1-dependent phosphorylation of its activation loop (Kobayashi & Cohen, 1999; Kobayashi *et al*, 1999). A third site, called the turn motif, also in the tail, regulates the stability of Akt and protein kinase C (PKC) (Ikenoue *et al*, 2008; Facchinetti *et al*, 2008), but its phosphorylation has not been reported in Sgk3. Distinct from the activation of Akt by PIP_3_ or PI(3,4)P_2_, Sgk3 depends on the lipid phosphatidylinositol-3-phosphate (PI3P) for its activation (Virbasius *et al*, 2001). Sgk3 is recruited specifically to endosomes via its PI3P-binding phox homology (PX) domain (Tessier & Woodgett, 2006; Xing *et al*, 2004), which promotes phosphorylation of its activation loop and hydrophobic motif by PDK1 and mTORC2 respectively (Tessier & Woodgett, 2006; Bago *et al*, 2014, 2016). Two major pathways drive the production of PI3P on endosomes: downstream of phosphatidylinositol-3,4,5-trisphosphate (PIP_3_) production by class I PI3K at the plasma membrane, phosphatidylinositol-3,4-bisphosphate [PI(3,4)P_2_] is produced by the action of SH2-containing inositol 5’ phosphatase (SHIP2) (Liu *et al*, 2018; Goulden *et al*, 2019). Phosphatidylinositol-3-phosphate (PI3P) is generated either from PI(3,4)P_2_ by inositol polyphosphate 4-phosphatases (INPP4) (Ivetac *et al*, 2005) or, de novo, from phosphatidylinositol by the class III PI3K Vps34 (Backer, 2008). Both pathways have been shown to contribute to Sgk3 activation (Bago *et al*, 2014; Malik *et al*, 2018).

Sgk isoforms are transcriptionally induced by serum and glucocorticoid hormones (Webster *et al*, 1993) and are involved, among others, in the regulation of ion channels, transcription factors, enzymatic activities and have been implicated in hair growth defects and renal dysfunction (Lang *et al*, 2006). Recently, Sgk3 has been shown to play a role in different cancers in an Akt-independent manner (Gasser *et al*, 2014; Bago *et al*, 2016; Vasudevan *et al*, 2009; Wang *et al*, 2017), highlighting the importance of Sgk3 as an effector downstream of PI3K signaling. Despite its physiological and pathological relevance, however, relatively little is known about the precise mechanism of Sgk3 regulation. In this manuscript, we provide clear evidence for the allosteric activation of Sgk3 by PI3P, highlighting the role of the PX domain and PI3P in the regulation of Sgk3 enzymatic activity. Our findings have obvious implications for the spatial and temporal control of Sgk3 activity as well as for the design of Sgk3-targeted therapeutics.

## Results

### Sgk3 localizes specifically to PI3P-containing endosomes

The Sgk3 PX domain (residues 1-124) has previously been shown to bind to PI3P, and is required for localization at early endosomes (Xu *et al*, 2001; Chandra *et al*, 2019; Virbasius *et al*, 2001; Bago *et al*, 2014). Serum-starved HeLa cells transfected with mCherry-Sgk3 show a punctate endosomal membrane localization (Fig. 1A), which is lost upon 20 min treatment with the PI3K inhibitor wortmannin (Fig. 1B). The membrane localization of Sgk3 is strictly dependent on its PX domain, since a binding site mutant of Sgk3 (Sgk3^R90A^) fails to localize to endosomes in the same cells (Fig. 1C-D). This is further illustrated by a representative cross section intensity profile (Fig. 1E-F). These data confirm both the specificity of Sgk3 for PI3P *in vivo* and the endosomal localization of Sgk3 to endosomes.

**Figure 1.**
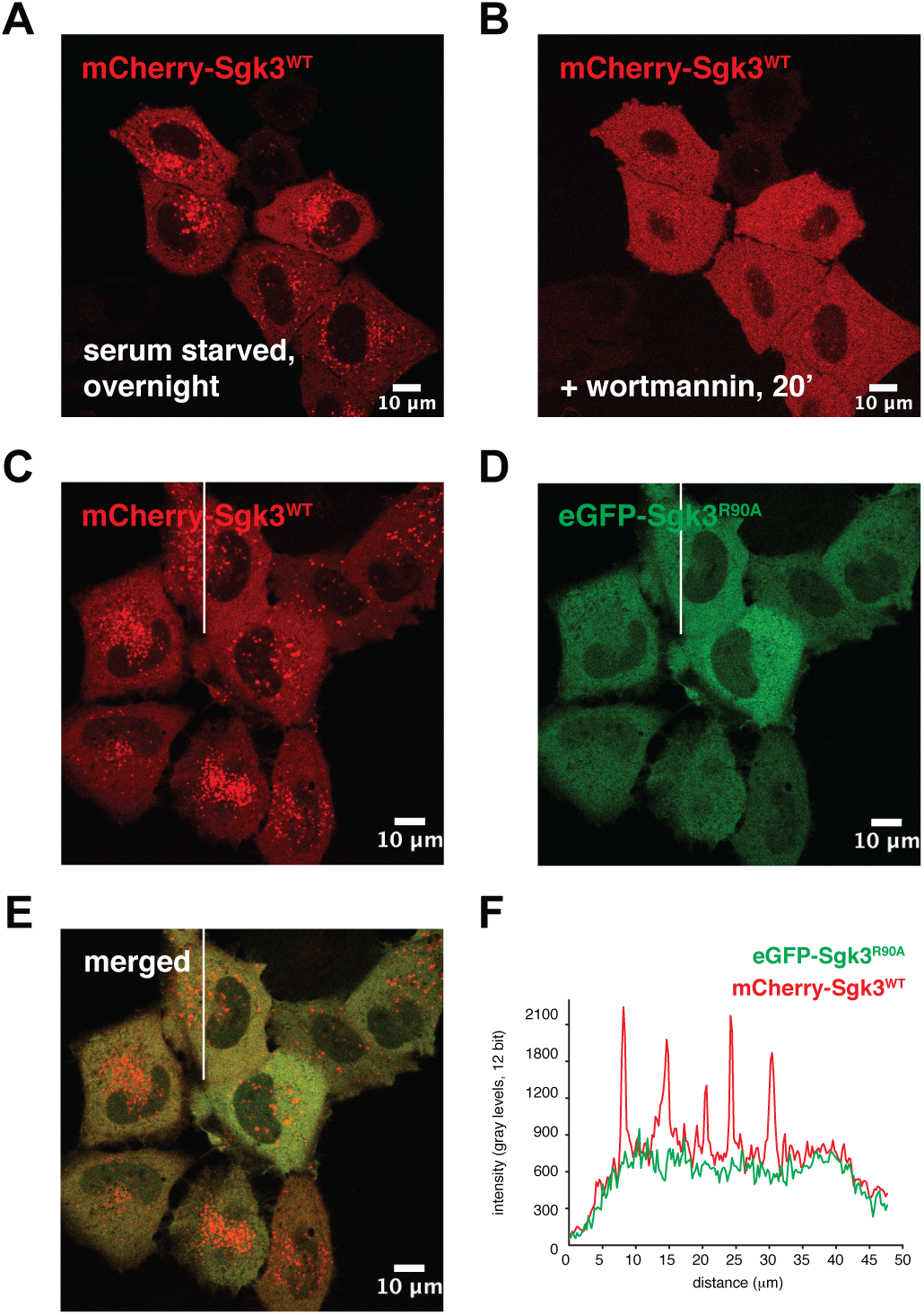
Sgk3 localizes specifically to PI3P-containing endosomes. **A)** HeLa cells were transfected with mCherry-Sgk3^WT^ and serum starved overnight. **B)** Starved cells were treated with 100 nM wortmannin for 20 min. **C-E)** Cells were co-transfected with mCherry-Sgk3^WT^ and EGFP-Sgk3^R90A^ mutant. **F)** Fluorescence intensity profile of mCherry Sgk3^WT^ and EGFP-Sgk3^R90A^ along the white line shown in C-E.

### *In vitro* characterization of recombinant Sgk3

The phosphorylation state of recombinant human Sgk3 co-expressed with PDK1 and purified from baculovirus-infected insect cells was characterized by tandem mass spectrometry (Supplementary Figure 1A). Sub-stoichiometric (∼20%) activation loop (Thr320) phosphorylation was observed; however, phosphorylated peptides were not observed for either the turn motif (Ser461) or the hydrophobic motif (Ser486), despite purification with phosphatase inhibitors known to preserve phosphorylation at these sites in Akt1 (manuscript in revision). Turn motif phosphorylation has previously been reported for Sgk1 (Chen *et al*, 2009) and other AGC kinases, in particular Akt and PKC, in which it plays an important role in regulating protein stability and degradation (Oh *et al*, 2010; Facchinetti *et al*, 2008; Ikenoue *et al*, 2008; Hauge *et al*, 2007). Since both the turn motif and the basic patch to which it binds on the kinase domain are conserved, it seemed logical to assume that Sgk3 should also be modified at this site. Insects, however, do not express an obvious homolog of Sgk3, raising the possibility that the kinase responsible for turn motif phosphorylation of Sgk3 is also not present in insect cells. We therefore expressed human Sgk3 in HEK293 cells and characterized its phosphorylation state by tandem mass spectrometry. Surprisingly, phosphorylated peptides were again not observed for either the turn motif or hydrophobic motif (Supplementary Figure 1B). It is therefore an open question as to whether Sgk3 is indeed modified on the turn motif and under what cellular conditions.

To prepare site-specifically phosphorylated Sgk3 for in vitro kinase assays, both wild-type Sgk3 (Sgk3^WT^) and kinase-dead Sgk3 (Sgk3^D286A^) were dephosphorylated with lambda phosphatase and re-phosphorylated with recombinant PDK1. Specific modification of Sgk3 with a single phosphate on its activation loop (Thr320) was confirmed by mass spectrometry (Supplementary Figure 1C-D). The purification of recombinant Sgk3 was challenging, due to solubility issues under certain purification conditions. However, careful optimization of the purification protocol enabled us to purify Sgk3 in a detergent-free buffer with pH and ionic strength, close to physiological values (see Materials and Methods). We performed size-exclusion chromatography coupled to multi-angle light scattering to determine its precise mass and monodispersity. Sgk3 exhibited a single peak with a measured mass of 57.3 kDa, in close agreement with its actual mass reported by mass spectrometry and a polydispersity value of 1.001 (Supplementary Figure 1E). To ensure that the Sgk3 used in subsequent assays was monodisperse and remained so for the duration of the assays, we measured the particle size distribution of Sgk3 by dynamic light scattering at 0 and 60 min post-freeze-thaw under kinase assay buffer conditions. Sgk3 exhibited near identical mass distributions centered around a particle size of 3 nm at 0 and 60 min at 20°C (Supplementary Figure 1F).

### Sgk3 is specifically and allosterically activated by PI3P

Previous studies have shown that loss of PI3P binding decreases phosphorylation of Sgk3 on its activation loop and hydrophobic motif (Tessier & Woodgett, 2006), as well as downstream substrate phosphorylation (Bago *et al*, 2016). However, it is not known whether PI3P binding serves to allosterically activate Sgk3 in addition to its known role in promoting phosphorylation of the activation loop and hydrophobic motif (Bago *et al*, 2014). Previous studies have established that the closely-related kinase Akt depends on the binding of PI(3,4,5)P_3_ or PI(3,4)P_2_ for full activity (Ebner *et al*, 2017; Lučić *et al*, 2018). We therefore investigated whether PI3P is simply a membrane anchor to target Sgk3 to endosomes, or whether PI3P also allosterically governs Sgk3 activity.

Having established that our recombinant Sgk3 is well-behaved during our assays, we assessed the influence of PI3P on kinase activity. Dephosphorylated Sgk3 exhibited undetectable kinase activity in vitro and could not be stimulated with PI3P (Supplementary Figure 1G). To test whether PI3P allosterically activates Sgk3, we bound PDK1-phosphorylated Sgk3^WT^ to liposomes with increasing PI3P concentrations and assayed its kinase activity against Crosstide, a glycogen synthase kinase 3β (GSK3β)-derived peptide substrate of Sgk3 (Dai *et al*), using a radiometric kinase assay (Hastie *et al*, 2006). We observed robust, PI3P-dependent activation of Sgk3, which correlated with the fraction of protein bound to the liposomes (Fig. 2A). Both the activity and binding curves can be fitted with a one site binding model characterized by nearly identical binding affinities of 24.6±10.8 µM and 29.3±8.0 µM, respectively. No signal was observed in the control without Crosstide, thereby ruling out autophosphorylation of Sgk3. Basal kinase activity against Crosstide seen in the absence of PI3P can be explained by the high degree of activation loop phosphorylation (Fig. S1C). To confirm that the observed activity comes exclusively from Sgk3, we mutated D286 in the catalytic loop to alanine to create a kinase-dead Sgk3. While this protein shows almost identical PI3P binding to Sgk3^WT^, neither basal nor PI3P-stimulated Crosstide phosphorylation was observed (Fig. 2B).

**Figure 2.**
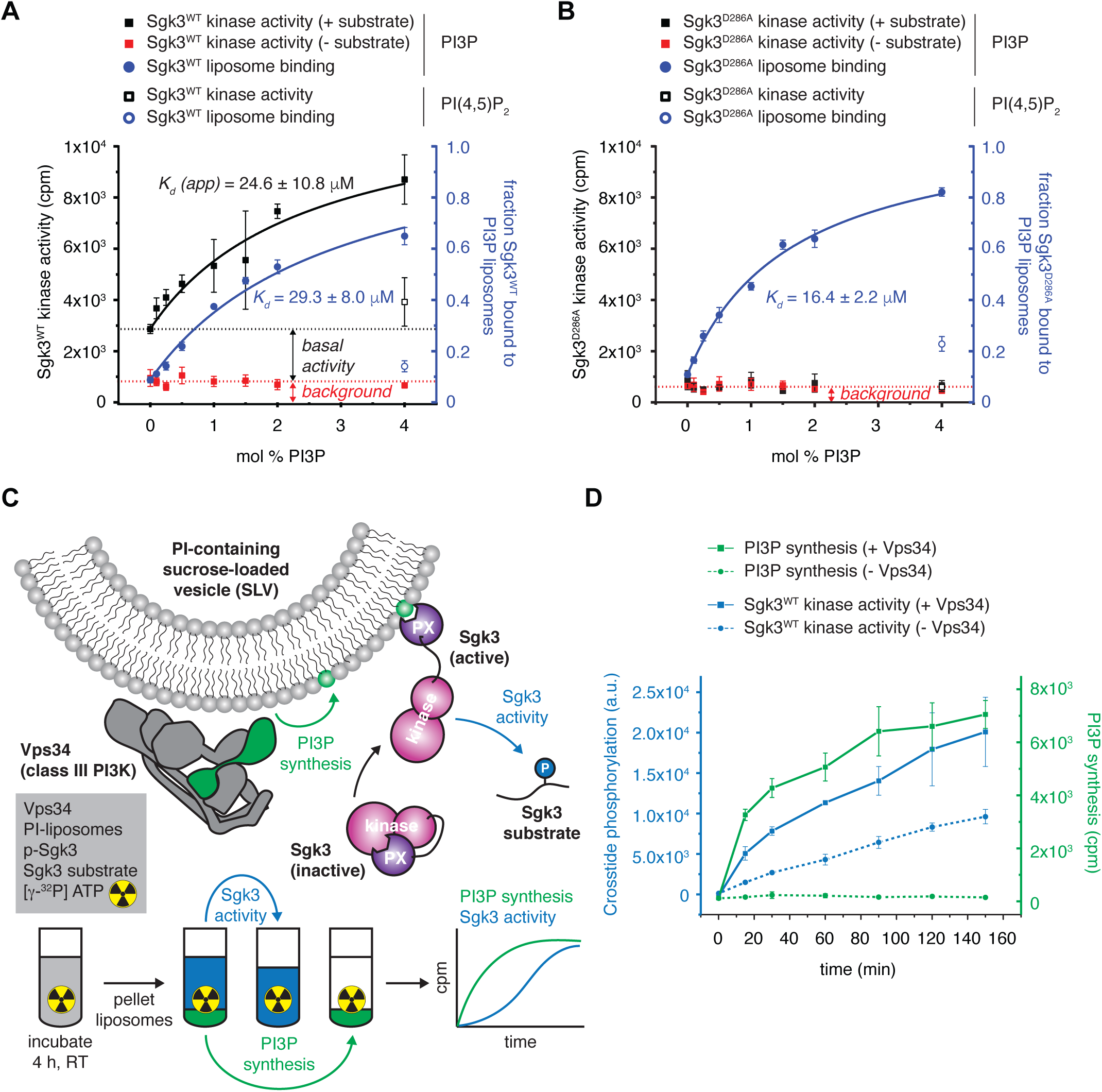
Sgk3 is specifically and allosterically activated by PI3P. **A)** Sgk3^WT^ kinase activity was assayed in the presence (closed black square) or absence (closed red square) of 100 µM Crosstide substrate with 0-4 mol % PI3P liposomes, or with 4 mol % PI(4,5)P_2_ liposomes (open black square) as a negative control. In parallel, binding of Sgk3 to 0-4 % PI3P liposomes (full blue circles) or 4% PI(4,5)P_2_ (open blue circle) as a negative control was measured. Dotted black line indicates basal Sgk3^WT^ kinase activity of unbound protein, dotted red line indicates no-substrate control. Kinase activity was read out radiometrically by liquid scintillation and liposome binding by Coomassie densitometry. Error bars are the standard deviation of three independent experiments. **B)** Same as A) with Sgk3^D286A^ kinase-dead mutant. **C)** Schematic of Vps34-Sgk3 coupled kinase assay. Phosphorylation of Crosstide was assayed by phosphorimaging. PI3P synthesis was measured by liquid scintillation. **D)** Vps34-Sgk3 coupled kinase assay. Crosstide phosphorylation was measured in the presence (full blue line) or absence (dashed blue line) of 50 nM Vps34. At the same time, PI3P synthesis was measured in the presence (full green line) or absence (dashed green line) of 50 nM Vps34. Error bars are the standard deviation of three independent experiments.

We next reconstituted Sgk3 activation by PI3P in vitro using a coupled Vps34-Sgk3 kinase activity assay. Vps34 is the only class III PI3K expressed in mammalian cells and generates PI3P from PI on endosomes and other endomembrane structures, including autophagosomes (Hurley & Young, 2017; Backer, 2016). We prepared liposomes containing 10 mol % PI as a substrate for Vps34 and monitored both Vps34 and Sgk3 activity simultaneously in a radiometric kinase assay (Fig. 2C). We could detect a time-dependent production of PI3P in the liposomes and the consequent activation of Sgk3 as a function of increasing PI3P (Fig. 2D). This demonstrates that Sgk3 is allosterically activated by the product of class III PI3K activity, highlighting that full Sgk3 activity requires membranes enriched in PI3P.

### HDX-MS reveals conformational changes in Sgk3 upon membrane binding

Having demonstrated that Sgk3 is allosterically activated by PI3P, we next set out to investigate the structural basis of PI3P-mediated activation. In previous studies on Akt, we have shown that PI(3,4,5)P_3_ or PI(3,4)P_2_ binding relieves pleckstrin homology (PH) domain-induced steric occlusion of the kinase domain catalytic cleft, leading to Akt activation (Ebner *et al*, 2017; Lučić *et al*, 2018). To verify whether this is also the case for Sgk3, we performed hydrogen-deuterium exchange mass spectrometry (HDX-MS) on full-length Sgk3 and Sgk3 PX domain in the presence of vesicles containing 0 or 5 mol % PI3P. Binding of the isolated PX domain to PI3P vesicles led to dramatically lower exchange rates for peptides across the domain, including the PI3P binding site, indicating that PI3P binding is captured by HDX-MS analysis (Fig. 3A). We next compared the HDX rates between full-length Sgk3^WT^ and the isolated PX domain to determine whether they interact with each other. We observed dramatic deprotection of the PX domain in the absence of the kinase domain of Sgk3 (Fig. 3B), indicating that the PX domain is sequestered in an intramolecular interaction in the full-length protein. Finally, we monitored HDX changes in full-length Sgk3 upon PI3P binding. Protection of the PX domain upon membrane binding was also observed, accompanied by deprotection in several areas of the kinase domain. These sites form a contiguous network starting with β5, β7 and β8 strands in the N-lobe and propagating to the C-lobe via the αG and αI helices (Fig. 3C). Additional deprotection was observed in residues 468-476 localized in the C-terminal tail. It was not possible to delineate a clear surface of interaction between the PX and kinase domains from the changes in exchange rates in the kinase domain, but the data, taken together, provide unambiguous evidence for an intramolecular PX-kinase domain interaction and a large conformational change upon PI3P binding.

**Figure 3.**
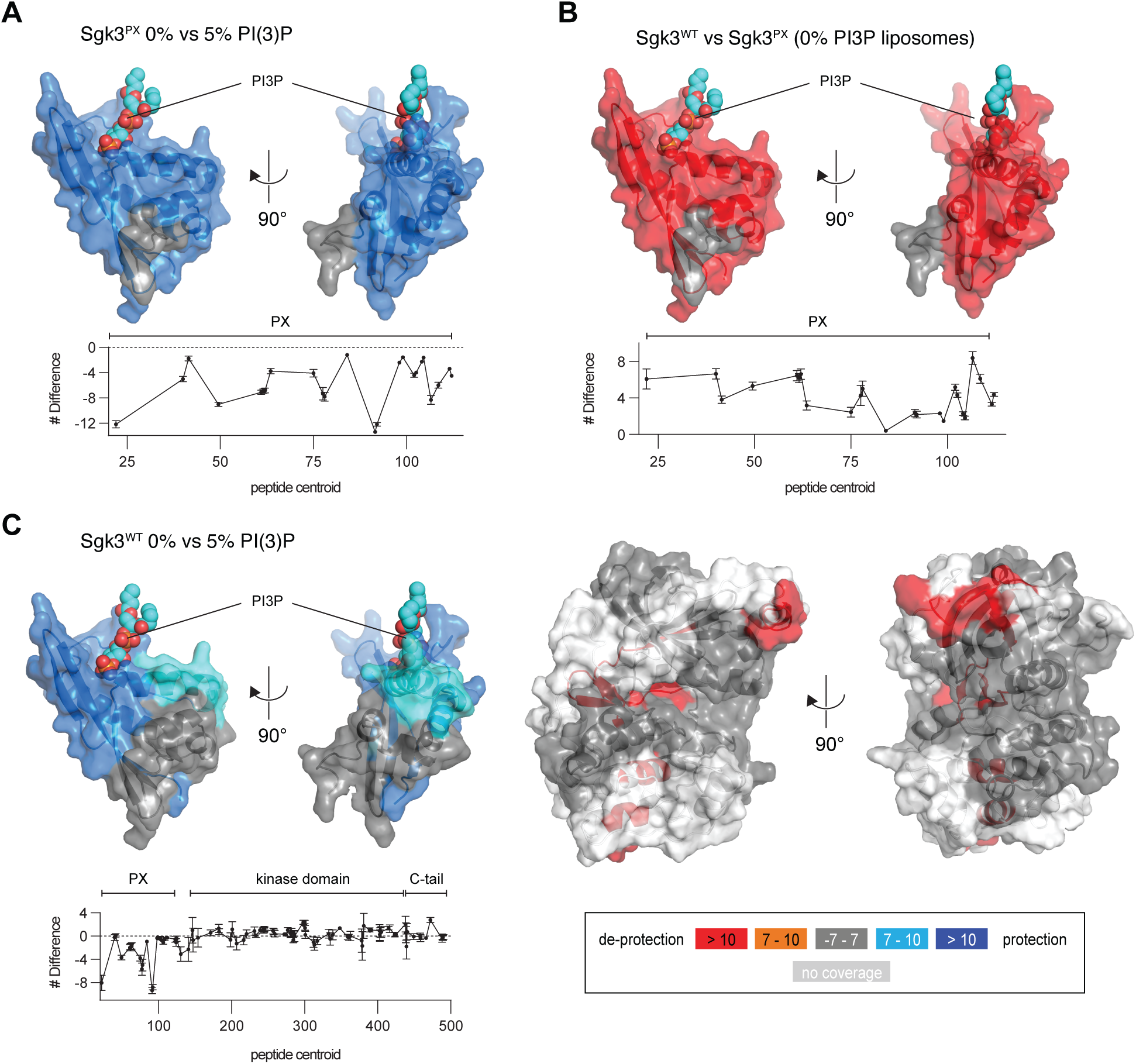
Sgk3 is autoinhibited by its PX domain. **A)** Comparison of deuterium incorporation in Sgk3 PX domain with 0% vs 5% PI3P liposomes. Plot: difference in number of deuterons incorporated as a function of peptide centroid over the PX domain. Negative numbers indicate protection (blue). Peptides showing significant deuterium exchange differences (>7%, >0.5 Da, and p<0.01) were mapped on the structure of the human Sgk3 PX domain (PDB:6EDX). The binding site for PI3P is indicated by superimposing the structure of the PX domain of p40phox in complex with PI3P (PDB: 1H6H) and rendering the PI3P in spheres. **B)** Comparison of deuterium incorporation in Sgk3^WT^ vs. Sgk3 PX domain in the presence of 0% PI3P liposomes (unbound). Plot: difference in number of deuterons incorporated as a function of peptide centroid over the PX domain. Positive numbers indicate domain exposure (red). **C)** Comparison of deuterium incorporation in Sgk3^WT^ unbound (0% PI3P) vs bound (5% PI3P) for the PX domain (left) and the kinase domain (right). A homology model of Sgk3 kinase domain (Rosetta) (Song *et al*) was used to map the changes. Color coding expressing the magnitude of changes in deuterium incorporation is on bottom right. All measurements were done in triplicates, error bars are the standard deviations.

### Autoinhibition of Sgk3 impairs binding to PI3P

Many AGC kinases that are activated by lipid second messengers exhibit a closed conformation of their autoinhibited state in which the binding site for the lipid headgroup is sequestered in an intramolecular interaction. To investigate whether the autoinhibited conformation of Sgk3 might also sequester the PI3P-binding site of its PX domain in intramolecular contacts, we compared the binding affinity for PI3P-containing liposomes of full-length Sgk3 and its isolated PX domain. To ensure identical conditions, we incubated both proteins with PI3P liposomes in a single reaction, pelleted the liposomes, resolved the proteins in the supernatant and pellet fraction by SDS-PAGE, and quantified their abundance by Coomassie stain densitometry. To compensate for the lower staining sensitivity of the PX domain, we used an N-terminally FITC-labelled version of this construct and quantified its fluorescence intensity in the gel before Coomassie staining. We observed an approximately 5-fold higher affinity to PI3P of the isolated PX domain over the full-length protein (Fig. 4A), confirming that the PI3P binding site is sequestered by the kinase domain in the autoinhibited state.

**Figure 4.**
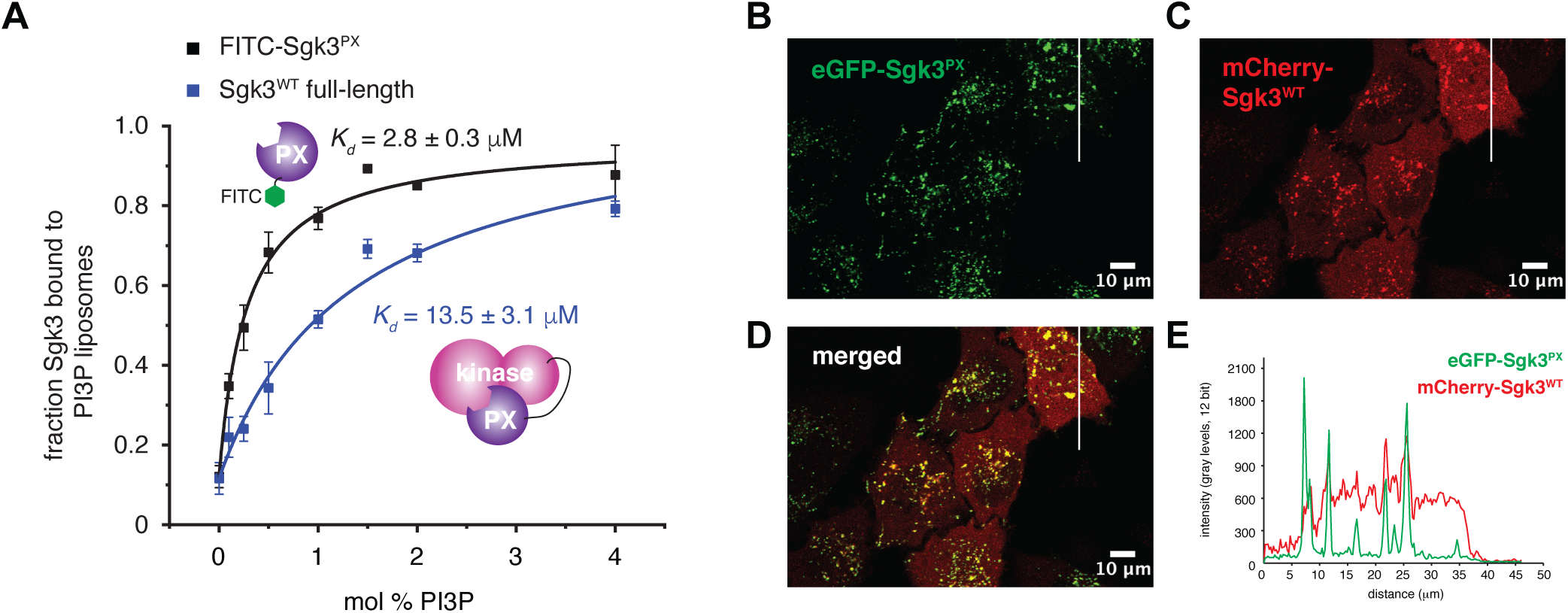
Autoinhibition of Sgk3 impairs binding to PI3P. **A)** Binding of Sgk3^WT^ full-length (blue squares) and FITC-labelled Sgk3 PX domain (black squares) to 0-4 mol % PI3P liposomes was monitored in single reaction using fluorescence (FITX-PX) or Coomassie densitometry (full-length Sgk3) in 15% SDS-PAGE gels. Curves were fitted with a one binding site model. Error bars are the standard deviations of three independent experiments. **B-D)** Distribution of EGFP-PX domain (B) or mCherry-Sgk3^WT^ (full-length) (C) in serum-starved HeLa cells. **E)** Quantification of fluorescence intensity of both constructs along the white line shown in images B-D.

To substantiate this finding in vivo, we expressed mCherry-Sgk3^WT^ and eGFP-Sgk3^PX^ in HeLa cells and imaged their localization in serum-starved cells by confocal microscopy. While serum starvation of the cells overnight did not reduce PI3P levels sufficiently to dissociate Sgk3 from endosomes (Fig. 1A), it was clear that the isolated Sgk3 PX domain associates almost exclusively with endosomes (Fig. 4B), while full-length Sgk3 exhibits a predominantly cytosolic distribution (Fig. 4C). A representative cross-section intensity trace for both channels reveals that the bulk of full length Sgk3 is uniformly distributed in the cytoplasm of cells, while the PX domain is exclusively found on endosomes (Figure 4D-E). These findings confirm that the PI3P binding pocket is sequestered in the autoinhibited conformation of Sgk3.

## Discussion

Sgk3 is dependent on both PI3P and phosphorylation for its activity. We have shown here that Sgk3 exhibits negligible activity in the absence of activation loop phosphorylation and that PI3P significantly increases the basal activity of pre-phosphorylated Sgk3. Like other AGC kinases regulated by phospholipids, including Akt and PKC, the PI3P binding pocket of the PX domain of Sgk3 is sequestered in an autoinhibitory, intramolecular conformation. This both necessitates PI3P for activity and simultaneously raises the threshold level of PI3P that must be reached in the cell in order to activate Sgk3. The structure of Akt1 reveals an intramolecular interaction between its PH and kinase domains that obscures the PIP_3_ binding site (manuscript in revision), a feature which impairs its binding to PIP_3_ in vitro and in vivo (Ebner *et al*, 2017). Similarly, the diacylglycerol (DAG) binding site of the C1b domain of PKCβII is also blocked by intramolecular contacts that impair its membrane translocation in response to phorbol ester treatment (Leonard *et al*, 2011). More generally, the regulatory C1 domains of all PKC isozymes, as well as those of the PKD family, are engaged in autoinhibitory intramolecular interactions that impair their binding to membranes (Oancea & Meyer, 1998; Lučić *et al*, 2016; Elsner *et al*, 2019). Mechanistically, Sgk3 exhibits many of the features previously described for Akt, though given the structural divergence between the PX domain of Sgk3 and the PH domain of Akt, the atomistic details of autoinhibition are likely to be different. Sgk3 is the only Sgk to comprise a full PX domain, although Sgk1 has been shown to possess a short ‘PX-like’ domain in its N-terminus that mediates lipid binding (Pao *et al*, 2007). Importantly, activation loop phosphorylation by PDK1 is insufficient to override the requirement for PI3P for full Sgk3 activity, a property also shared by Akt (manuscript in revision).

Whilst not the major focus of this study, it is worth noting that turn motif phosphorylation of Sgk3 could not be observed in Sgk3 expressed either in insect or human cells. This finding was unexpected, given the prominent role of turn motif phosphorylation in other AGC kinases. Phosphorylation of the turn motif helps to anchor the C-terminal tail of the kinase domain and increase the affinity of the phosphorylated hydrophobic motif to its cognate hydrophobic pocket in the N-lobe via local concentration, which leads to the overall stabilization of the kinase fold (Hauge *et al*, 2007). While turn motif phosphorylation has been reported in all PKC isoforms, Akt, Sgk1, and PKN (Hornbeck *et al*, 2004), whether it plays a role in Sgk3 regulation will require further investigation. Finally, it should also be noted that while hydrophobic motif phosphorylation has been reported to increase the catalytic activity of Sgk3 (Bago *et al*, 2014), our study does not address its contribution to Sgk3 activation.

Evidence suggests that PI3P binding by Sgk3 also promotes PDK1-dependent phosphorylation of its activation loop. This mechanism of activation loop exposure upon membrane binding is well established for Akt (Alessi *et al*, 1997; Stokoe *et al*, 1997). The requirement of PI3P for Sgk3 activity is therefore three-fold: in the first place, it is required to target Sgk3 to membranes where it can encounter active PDK1; secondly, PI3P binding triggers a conformational change that renders it a substrate for PDK1; thirdly, and importantly, it is required to prevent autoinhibition of the kinase domain by the PX domain. Studies on Akt have also implicated ATP in stabilizing a phosphatase-resistant conformation of the kinase domain (Lu *et al*, 2015; Lin *et al*, 2012; Chan *et al*, 2011), a mechanism which we recently showed to be dependent on binding to membrane-embedded PIP_3_ (Lučić *et al*, 2018). While such a mechanism has not been explicitly shown for Sgk3, the homology between the kinase domains and the mechanisms by which Akt and Sgk3 are activated suggests that this is also likely to be the case for Sgk3 bound to PI3P-rich endosomes.

Our findings suggest that PI3P and phosphorylation are both required for Sgk3 activity, which makes Sgk3 a coincidence detector of both PI3P and PI3K signaling. According to this model, PI3P-decorated endosomes lacking PDK1 would not serve as signaling hubs for Sgk3. In this respect, it is worth noting that PDK1 does not bind to, nor can it be activated by, PI3P. PDK1 binds with almost equal affinity to both PI(3,4,5)P_3_ (1.6 nM) and PI(3,4)P_2_ (5.2 nM) (Currie *et al*, 1999), which targets it to both the plasma membrane, which is abundant in PIP_3_, and endomembranes, which are decorated with PI(3,4)P_2_ and heavily implicated in Akt signaling (Liu *et al*, 2018; Ebner *et al*, 2017; Schenck *et al*, 2008; Braccini *et al*, 2015). PI3P is generally not observed at the plasma membrane. Presumably, PDK1 and Sgk3 cooperate to drive Sgk3 signaling on endosomes containing both PI(3,4)P_2_ and PI3P (Figure 5). Indeed, PDK1 has also been reported to associate with endosomes dependent on Sgk3 hydrophobic motif phosphorylation, where it inhibits ligand-induced degradation of the chemokine receptor CXCR4 (Slagsvold *et al*, 2006). Sgk3 activity could therefore be restricted to a subpopulation of endosomes with a specific phosphoinositide composition. Interestingly, a recent large scale analysis of human PX domains revealed the presence of a secondary lipid binding site in the PX domain of Sgk3 which exhibits low-affinity PI(3,4)P_2_ binding (Chandra *et al*, 2019). The functional importance of the second lipid-binding site is still unclear, though one could easily imagine that it might serve to either fine-tune the lipid binding specificity or increase binding affinity via an avidity effect (Carlton & Cullen, 2005).

**Figure 5.**
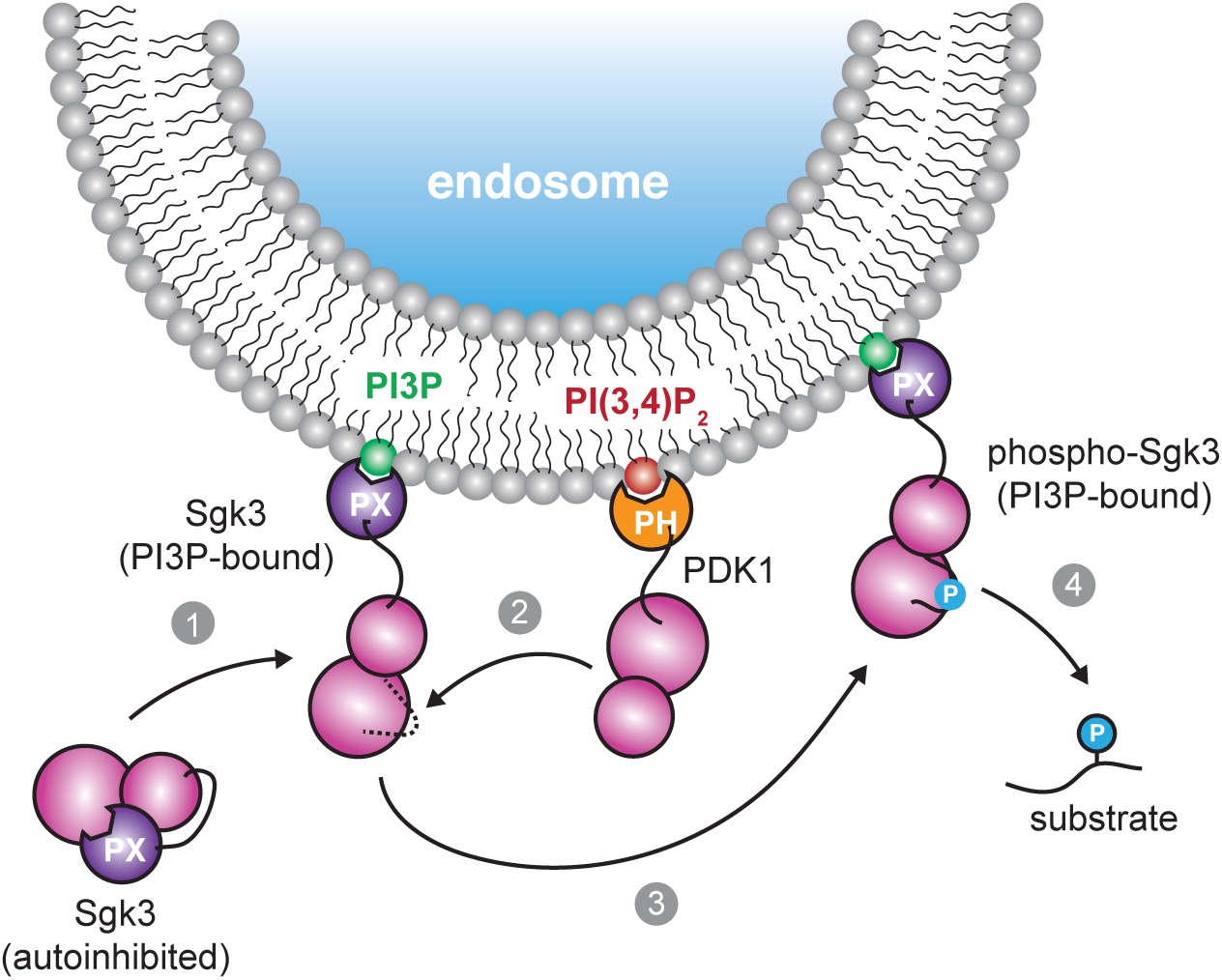
Model for the allosteric activation of Sgk3 by PI3P. Cytosolic Sgk3 is allosterically inhibited by its PX domain, which is relieved by PI3P binding (1). This is followed by phosphorylation of the Sgk3 activation loop by PDK1 as a consequence of its association with endosomal PI(3,4)P_2_ (2). Phosphorylation activates Sgk3 (3), thereby permitting downstream substrate phosphorylation (4).

A similar model for PI(3,4)P_2_-driven Akt activity on endomembranes has recently been proposed to explain the transduction of growth factor signaling by Akt at subcellular locations distal to the plasma membrane. In this model, both PDK1 and Akt are targeted to PI(3,4)P_2_-containing endomembranes, where they cooperate to drive Akt signaling in the interior of the cell. Quantitative lipid imaging has revealed that endosomal pools of PI(3,4)P_2_ are signaling hubs for Akt (Liu *et al*, 2018). Like Sgk3, Akt activity depends strictly on phosphorylation and the binding of its PH domain to either PIP_3_ or PI(3,4)P_2_, thereby confining its activity to membranes enriched in these signaling lipids (Ebner *et al*, 2017; Lučić *et al*, 2018; Siess & Leonard, 2019).

Such a model of localized Sgk3 activity may provide an explanation for the reported overlap in Akt and Sgk3 substrate specificity (Kobayashi *et al*, 1999; Tessier & Woodgett, 2006; Brunet *et al*, 2001; Murray *et al*, 2005). Notwithstanding the similarity in their kinase domains, it is clear that their activity profiles would overlap in such a model, since the PI(3,4)P_2_ that drives PDK1 and Akt activation must be present in membranes that support PI3P- and PDK1-mediated Sgk3 activation. Indeed, a recent phosphoproteomic study identified several Sgk3-specific substrates localized on endosomes (Malik *et al*, 2019). The reported role of Sgk3 in driving human cancers in the absence of Akt hyperactivation is then also reconcilable with this model and reinforces the potential value of Sgk3 as a therapeutic target either as a monotherapy or in combination with Akt inhibitors.

The chemical toolkit for Sgk3 inhibition is still limited due to similarities with other AGC kinases, such as Akt, and indeed with Sgk1 and Sgk2. However, inhibitor 14h has been reported to target all three Sgk isoforms without inhibiting Akt (Bago *et al*, 2016) and, more recently, an Sgk3-specific PROTAC degrader has been described (Tovell *et al*, 2019). Much effort has been aimed at developing allosteric inhibitors of Akt that specifically target the autoinhibited conformation of the kinase (Wu *et al*, 2010; Lapierre *et al*, 2016; Weisner *et al*, 2019), although despite high oral bioavailability and specificity, none have yet made it to phase III clinical trials (Landel *et al*, 2020). Nevertheless, the identification of an autoinhibitory intramolecular interface in Sgk3, analogous but not homologous to Akt, presents opportunities for the development of Sgk3-specific inhibitors that may have clinical value in the treatment of cancers in which Sgk3 signaling is observed to be upregulated. Such studies will undoubtedly benefit from the high-resolution structure determination of full-length, autoinhibited Sgk3.

## Acknowledgements

Intact mass analyses and phosphorylation mapping experiments were performed by Dorothea Anrather and Thomas Gossenreiter from the Mass Spectrometry Facility at the Max Perutz Labs using instruments of the Vienna BioCenter Core Facilities (VBCF). We thank Justyna Sawa-Makarska and Sascha Martens (Max Perutz Labs) for their generous gift of baculovirus for the expression of the Vps34 complex. This work was supported by the Austrian Science Fund (FWF) grants P28135, P30584 and P33066 to T.L. L.T. was supported by a Hertha Firnberg Postdoctoral Fellowship from the Austrian Science Fund (FWF T915). J.E.B. is supported by a Michael Smith Foundation for Health Research (MSFHR) Scholar award (17686), and an operating grant from the Cancer Research Society (CRS-24368).

## Author Contributions

D.P. purified all proteins and performed all biochemical experiments. L.T. performed all fluorescence microscopy experiments. K.F. and J.B. performed all HDX-MS experiments and analyzed the corresponding data. T.L. conceived the project.

## Materials and methods

### Fluorescence microscopy

For localization studies of SGK3 in live cells, HeLa cells were co-transfected with mCherry-SGK3 (wt) and either eGFP-SGK3^PX^ (residues 1-124) or eGFP-SGK3^R90A^. Cells were transfected using Turbofect transfection reagent according to the manufacturer’s protocol. Cells were serum starved overnight and changed into HBSS for live imaging. Images were acquired 24 hours after transfection on a Zeiss LSM710 or LSM 700 confocal microscope equipped with a Zeiss 63x oil immersion objective (NA 1.4) and 488 and 561 nm lasers. Image processing and analysis were done using ImageJ 1.51j. All adjustments were applied uniformly across the entire image; no nonlinear adjustments were used.

### Protein expression and purification

Sgk3 PX domain (residues 1-124) was cloned into pHis-parallel bacterial expression vector and transformed into *E. c*oli BL21 STAR electrocompetent cells. For large-scale expression, the cells were grown in LB medium containing 100 µg/ml ampicillin at 37°C/180 RPM shaking, until OD_600_ reached 0.6. The temperature was lowered to 20°C, expression was induced by the addition of 0.1 mM IPTG and cells were grown overnight. Cells were harvested by centrifugation at 4000g/20 min, the pellet was flash-frozen in LN2 and stored at −80°C for further use. Pellet corresponding to 1 l of culture was resuspended in 100 ml PX lysis buffer: 50 mM Tris pH 7.5, 150 mM NaCl,

20 mM imidazole, 1% glycerol, 1 mM TCEP, 1 mM PMSF, 1 mM benzamidine hydrochloride, 2 mM MgCl_2_, 1x protease inhibitor cocktail (made in-house: 100 µM bestatin, 14 µM E-64, 10 µM pepstatin A, and 1 µM phosphoramidon), 5 µl Denarase. The cell suspension was sonicated on ice, spun at 38 724 g, 4°C for 30 min. The soluble fraction was loaded on a HisTrap FF 5 ml column (Cytiva) equilibrated in NiA buffer: 50 mM Tris pH 7.5, 150 mM Nacl, 20 mM imidazole, 1% glycerol, 1 mM TCEP. The column was washed with 20 column volumes (CVs) of NiA, 10 CVs of high salt wash buffer (NiA + 500 mM NaCl), an additional 10 CVs of NiA, and the bound protein eluted with 0.02-1 M imidazole gradient over 10 CVs. Fractions containing His-tagged PX domain were pooled and diluted with SA buffer in (50 mM Hepes pH 8.0, 1% glycerol, 1 mM TCEP). The His tag was cleaved with TEV protease (made in-house) overnight at 4°C. The cleaved protein was loaded on HiTrap SP HP 1 ml cation exchange column (Cytiva) equilibrated in SA buffer. The column was washed with 10 CVs of SA and eluted with a 0-1 M NaCl gradient over 20 CVs. Fractions containing the PX domain were diluted with SA to adjust NaCl concentration to ∼150 mM and were either used for Sortase labeling (see below) or concentrated to 20 mg/ml, snap-frozen in LN2 and stored at −80°C for further use.

Purified PX domain was specifically labeled on its N-terminal glycine with a FITC-LPETGG peptide (Genscript) in a Sortase-mediated reaction according to the protocol described in (Theile *et al*, 2013). Briefly, the PX domain was incubated with a 10-fold excess of FITC-LPETGG peptide in the presence of His-tagged Sortase A (prepared in-house) and 5 mM CaCl_2_ at 4°C overnight. Sortase was removed by NiNTA affinity chromatography and the labeled PX domain purified from free label by size exclusion chromatography on a Superdex 75 10/300 column equilibrated in 20 mM Tris pH 8, 150 mM NaCl, 1% glycerol, 20 mM imidazole, 1 mM TCEP. Fractions containing FITC-labeled PX domain were pooled, concentrated to 0.8 mg/ml, snap-frozen in LN2 and stored at −80°C.

The human Sgk3 gene was codon optimized for expression in insect cells (Genscript) and fused to the C-terminus of His_10_/StrepII-tagged eGFP (A206K monomeric variant) to aid expression of soluble Sgk3. Both wild-type and kinase-dead (D286A) Sgk3 were expressed in baculvirus-infected Sf9 insect cells grown in ESF 921 medium (Expression Systems) at 27°C. Cells were harvested 4-5 days post-infection. Wild-type and D286A constructs were purified under identical conditions. Briefly, the pellet of 0.5 l Sf9 cells was resuspended in 50 ml Sgk3 lysis buffer: 50 mM Hepes pH 7.4, 50 mM NaCl, 1% glycerol, 1 mM TCEP, 1 mM PMSF, 1 mM benzamidine, 1 mM MnCl_2_, 0.5 mM EDTA, 1x protease inhibitor cocktail, 0.1% CHAPS, 5 ul Denarase, 10 mM DTT, 1x protease inhibitor cocktail, 5 µl Denarase, 80 nM lambda phosphatase (made in-house). The lysate was passed through Dounce homogenizer several times and spun at 38 724 g (18 000 RPM), 4°C for 30 min. The soluble fraction was loaded onto a StrepTrap HP 5 ml column (Cytiva) equilibrated in 50 mM Hepes 7.4, 50 mM NaCl, 1% glycerol, 1 mM TCEP. The column was washed with 10 CVs of binding buffer and the protein eluted with 2.5 mM *d*-desthiobiotin in binding buffer. Fractions containing EGFP-Sgk3 were pooled and dephosphorylated with lambda phosphatase in 50 mM Hepes pH 7.4, 150 mM NaCl, 1% glycerol, 10 mM DTT, 1 mM MnCl_2_, 0.5 mM EDTA, 0.5 µM lambda phosphatase (made in-house). Dephosphorylation was carried out for 30 min at room temperature and 4°C overnight together with TEV cleavage of the His/StrepII tag. To separate His-tagged lambda phosphatase from Sgk3, the reaction mixture was loaded on HisTrap FF 1 ml column equilibrated in 20 mM Tris pH 7.4, 150 mM NaCl, 1% glycerol, 1 mM TCEP. Dephosphorylated and cleaved Sgk3 was washed from the column with 20 mM imidazole, while lambda phosphatase remained bound. Sgk3 was then site-specifically in vitro re-phosphorylated by GST-PDK1 under the following conditions: 50 mM Tris pH 7.4, 150 mM NaCl, 1% glycerol, 1 mM TCEP, 1 mM ATP, 2 mM MgSO_4_, 1:10 PDK1:Sgk3. The reaction was carried out at 4°C for 3 h, then stopped by addition of 10 mM EDTA. GST-PDK1 was separated by addition of glutathione-sepharose 4B beads (Cytiva) and incubation at 4°C overnight with gentle rotation. Supernatant containing rephosphorylated Sgk3 was concentrated in 30 kDa MWCO spin concentrator and injected onto Superdex 200 Increase 10/300 GL column (Cytiva) equilibrated in 20 mM Tris pH 7.4, 150 mM NaCl, 1% glycerol, 1 mM TCEP. Fractions containing Sgk3 were pooled, concentrated, snap-frozen in LN2 and stored at −80°C for further use.

Recombinant baculovirus encoding Vps34 complex I (Atg6, protein A-tagged Atg14, Vps15 and Vps34) (Sawa-Makarska *et al*, 2020) was used to infect 3 l of Sf9 insect cell culture. Cells were harvested 4-5 days post-infection. Pellets were lysed in 300 ml of Vps34 lysis buffer: 50 mM Tris pH 8.8, 300 mM Nacl, 1 mM DTT, 0.5 % CHAPS, 1 mM MgCl_2_, 0.5 mM PMSF, 1x protease inhibitor cocktail, 15 ul Denarase. The lysate was passed several times through a Dounce homogenizer and spun at 38 724 g (18 000 RPM), 4°C for 30 min. The soluble fraction was bound to IgG-Sepharose beads (Cytiva) for 1 h at 4°C with gentle rotation. Resin was collected in a gravity flow column and washed 3 x 50 ml wash buffer 1 (50 mM Tris pH 8, 300 mM NaCl, 1 mM DTT, 0.5% CHAPS, 0.5 mM PMSF), 2 x 50 ml wash buffer 2 (50 mM Tris pH 8, 300 mM NaCl, 1 mM DT, 2% glycerol, 0.25 % CHAPS). Finally, beads were resuspended in wash buffer 2 and bound protein complex cleaved with TEV at 4°C overnight. The supernatant containing the full Vps34 complex was concentrated and injected onto a Superdex 200 Increase 10/300 GL column equilibrated in 50 mM Tris pH 8, 150 mM NaCl, 1 mM TCEP, 2% glycerol. Fractions containing the complete complex of Atg6, Atg14, Vps15 and Vps34 were pooled, concentrated to 0.9 mg/ml, snap-frozen in LN2 and stored at −80°C for further use.

### Intact mass spectrometry

Intact protein samples were diluted in H_2_O and up to 100 ng protein were loaded on an XBridge Protein BEH C4 column (2.5 µm particle size, dimensions 2.1 mm X 150 mm; Waters) using a Dionex Ultimate 3000 HPLC system (Thermo Fisher Scientific) with a working temperature of 50 °C, 0.1% formic acid (FA) as solvent A, 80% acetonitrile, 0.08% FA as solvent B. Proteins were separated with a 6 min step gradient from 10 to 80% solvent B at a flow rate of 300 μL/min and analysed on a Synapt G2-Si coupled via a ZSpray ESI source (Waters). Data were recorded with MassLynx V 4.1 (Waters) and analyzed using the MaxEnt 1 process to reconstruct the uncharged average protein mass.

### Tandem mass spectrometry

The reduced protein was denatured in 4 M urea, 50 mM ammonium bicarbonate (ABC) and alkylated with 20 mM iodoacetamide for 30 min at room temperature in the dark. The sample was diluted with 50 mM ABC down to 0.5 M urea and then digested overnight using sequencing-grade endoproteinase GluC (Roche) at 25 °C. The digestion was stopped by adding trifluoroacetic acid to a final concentration of 1% and the peptides were desalted using custom-made C18 stagetips (Rappsilber *et al*, 2007). The peptides were separated on an Ultimate 3000 RSLC nano-flow chromatography system (Thermo Fisher Scientific), using a pre-column for sample loading (PepMapAcclaim C18, 2 cm × 0.1 mm, 5 μm) and a C18 analytical column (PepMapAcclaim C18, 50 cm × 0.75 mm, 2 μm, both Dionex-Thermo Fisher Scientific), applying a linear gradient from 2 to 35% solvent B (80% acetonitrile, 0.08% FA acid; solvent A: 0.1% FA) at a flow rate of 230 nl/min over 60 min. Eluting peptides were analysed on a Q Exactive HF or HF-X Orbitrap mass spectrometer equipped with a Proxeon nanospray source (Thermo Fisher Scientific), operated in data-dependent mode. Survey scans were obtained in a scan range of 375–1500 m/z, at a resolution of 60000 at 200 m/z and an AGC target value of 3E6. The 8 most intense ions were selected with an isolation width of 1.6 Da, fragmented in the HCD cell at 28% collision energy and the spectra recorded at a target value of 1E5 and a resolution of 30000. Peptides with a charge of +1 were excluded from fragmentation, the peptide match and exclude isotope features were enabled and selected precursors were dynamically excluded from repeated sampling. Raw data were processed using the MaxQuant software package (version 1.6.0.16, http://www.maxquant.org/) (Cox & Mann, 2008) and searched against a custom database containing the sequences of the Sgk3 constructs in the protein background of the expression system (either Spodoptera spp. protein sequences available in Uniprot plus the Bombyx mori reference proteome (www.uniprot.org) or the human proteome) plus the sequences of common contaminants. The search was performed with full trypsin specificity and a maximum of two missed cleavages, C-terminal cleavage to glutamate and aspartate for the GluC digests or unspecific cleavage for pepsin digests. Carbamidomethylation of cysteine residues was set as fixed, oxidation of methionine, protein N-terminal acetylation and phosphorylation of serine, threonine and tyrosine as variable modifications—all other parameters were set to default. Results were filtered at a false discovery rate of 10% at the peptide and protein level. Spectra of phosphorylated Sgk3 peptides were validated manually. All proteomics data have been deposited in the PRIDE repository (Perez-Riverol *et al*, 2019).

### Size exclusion chromatography with multiangle light scattering (SEC-MALS)

80 µl of Sgk3 at 0.9 mg/ml was injected onto Superdex 200 Increase 10/300 GL column (Cytiva) equilibrated in 20 mM Tris pH 8, 150 mM NaCl, 1% glycerol, 1 mM TCEP. Separation was carried out using HPLC Agilent Technologies 1260 infinity with a flow rate of 0.5 ml/min operated at room temperature. On-line light scattering of 690 nm laser was recorded on a miniDawn Treos detector (Wyatt Technology Corp.). Shodex RI-101 (Shodex) refractive index detector was used to measure protein concentration on-line. Data was analyzed using the ASTRA software (Wyatt Technology Corp.).

### Dynamic light scattering (DLS)

Autocorrelation curves of freshly thawed and centrifuged 7.5 µM Sgk3 in buffer containing 20 mM Hepes pH 7.4, 150 mM NaCl, 1 % glycerol, 1 mM TCEP, 1 mM ATP, 2 mM MgCl_2_, 100 µM Crosstide were recorded at 0 and 60 min of incubation at room temperature. Measurements were performed on a DynaPro NanoStar instrument (Wyatt Technology Corp.). Each measurement is an average of 10 acquisitions of 10 s each.

### Sucrose-loaded vesicles (SLVs)

Cholesterol, DOPC, DOPE, DOPS and liver PI were dissolved in chloroform, PI3P di C16 was dissolved in chloroform:methanol:water 1:2:0.8, PI(4,5)P_2_ was dissolved in chloroform:methanol:water 20:13:3. For liposomes with synthetic PI3P the lipids were mixed in the following molar ratio: 20% cholesterol, 30 % DOPC, 15% DOPS, 35% DOPE. For liposomes containing PI3P, DOPC was substituted with 0-4% PI3P. Liver PI liposomes contained 10% PI at the expense of DOPC. Lipid mixtures in borosilicate glass tubes were evaporated with nitrogen to form a thin film and then rehydrated by vortexing for 1 min with sucrose buffer (20 mM Hepes pH 7.4, 300 mM sucrose). Resuspended lipids were subjected to 4 cycles of freezing in LN2 and thawing in a sonicator bath. One volume of gel filtration buffer was added and liposomes were pelleted in Beckman Coulter Optima MAX-XP Ultracentrifuge in TLA 100 rotor operated at 50 000 rpm/20°C for 30 min. The supernatant was discarded, liposome pellets were resuspended in gel filtration buffer to a final lipid concentration of 1 mM and stored for a maximum of two days at 4°C before use.

### Radiometric kinase assays

Sgk3 kinase activity was monitored as the incorporation of radioactive phosphate from γ^32^P-ATP (Hartmann Analytic) into Crosstide (GRPRTSSFAEG), a validated Sgk3 substrate peptide. Sgk3 was mixed with PI3P-containing liposomes and incubated for 30 min at RT. 5 µl of liposomes-Sgk3 mixture was added to 5 ul of substrate master mix to obtain following conditions: 20 mM Tris pH 8, 150 mM NaCl, 1% glycerol, 1 mM TCEP, 50 nM Sgk3, 100 µM Crosstide, 1 mM ATP (spiked 1:10 with γ^32^P-ATP), 2 mM MgCl_2_. The reaction was left to proceed for 15 min at room temperature and stopped by addition of 2 µl of 0.5 M EDTA. 10 ul of the reaction was then spotted onto a nitrocellulose membrane (0.2 um pore size), washed 5 x 3 min with 75 mM H_3_PO_4_ and the washed membrane inserted into a 20 ml polyethylene scintillation vial containing 5 ml deionized H_2_O. Cerenkov radiation was measured in a Tri-Carb 4910 TR liquid scintillator (Perkin Elmer). Datapoints were fitted with Michaelis-Menten equation modified with a constant to reflect the basal enzymatic activity in 0% PI3P condition:

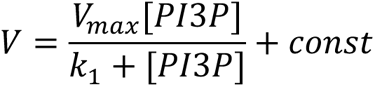

### Liposome pelleting assay

1 µM Sgk3^WT^ or Sgk3^PX^ was mixed 1:1 with liposomes, incubated at 20°C for 30 min and spun at 9800g for 30 min at 20°C. Supernatant was collected and the liposome pellet resuspended in gel filtration buffer. Equal volumes of supernatant and pellet fractions were separated by SDS-PAGE and quantified by Coomassie densitometry. In the comparative binding assay with full-length Sgk3 and Sgk3^PX^ in the same reaction, FITC-labelled PX domain was quantified by fluorescence at 448 nm. Binding curves were fitted with a one-site binding model.

### Vps34-Sgk3 coupled kinase assay

To monitor PI3P production by Vps34 and Crosstide phosphorylation by Sgk3 in a single reaction, the components were mixed at following concentrations: 50 nM Vps34 complex, 50 nM Sgk3, 500 µM Biotin-Crosstide, 0.5 mM ATP (spiked 1:10 with γ^32^P-ATP), 0.5 mM MgCl_2_, 2 mM MnCl_2_, 1 mM EGTA, 10% liver PI liposomes. The reaction was carried out at 25°C and gentle shaking. 15 µl samples were mixed with 5 µl 4x quench buffer (20 mM Tris pH 7.4, 150 mM NaCl, 10 mM EDTA, 10 mM EGTA, 1% glycerol, 1 mM TCEP) at indicated time points. The quenched reactions were spun for 5 min at 20800 g/20°C. The supernatant was mixed 1:1 with NeutrAvidin (Thermo Scientific) at 2 mg/ml in water to sequester the biotinylated Crosstide and 5 ul were spotted onto 0.45 um nitrocellulose membrane. The membrane was washed 5×3 min with TBS (20 mM Tris pH 7.6, 137 mM NaCl), briefly dried, exposed in storage phosphor screen overnight and imaged with an Amersham Typhoon phosphorimager. The intensity of the spots was quantified in ImageJ. The pellets containing liposomes with PI3^32^P were washed 4 times with wash buffer (20 mM Tris pH 7.4, 150 mM NaCl, 2.5 mM EDTA, 2.5 mM EGTA, 1% glycerol, 1 mM TCEP), transferred into PCR tube and inserted into 20 ml polyethylene scintillation vials containing 5 ml MilliQ water. PI3P synthesis was measured by liquid scintillation.

### HDX-MS

#### Lipid Vesicle Preparation

Lipid films containing 20% cholesterol, 25% DOPC, 15% DOPS, 35% DOPE, and 5% PI3P were dried and stored under argon for shipping. The dried films were resuspended to 5 mM in gel filtration buffer (20 mM Tris pH 7.4, 150 mM NaCl, 1% glycerol, 1 mM TCEP) by vortexing. The resuspended lipid vesicle solutions were sonicated for 10 minutes, subjected to ten freeze-thaw cycles, and sonicated again for 5 minutes. The vesicles were then snap-frozen in liquid nitrogen and stored at −80°C

#### HDX-MS sample preparation

HDX reactions were conducted in a final reaction volume of 25 μL with a final Sgk3 concentration of 13 μM (for both FL and PX Sgk3). Prior to the addition of deuterated solvent, 1.5 μL of FL or PX Sgk3 was allowed to incubate with either 2 μL of 5 mM 0% PI3P vesicles or 2 μL 5 mM 5% PI3P vesciles. After the two minute incubation period, μL D2O buffer (20mM pH 7 HEPES, 100mM NaCl, 0.5 mM TCEP, 94% D2O) was added with a final %D2O of 81.5% (v/v). The reaction was allowed to proceed for 3s, 30s, 300s, or 3000s at 18°C before being quenched with ice cold acidic quench buffer.

All conditions and timepoints were created and run in triplicate. Samples were flash frozen in liquid nitrogen and stored at −80°C until injection onto the ultra-performance liquid chromatography system for proteolytic cleavage, peptide separation, and injection onto a QTOF for mass analysis.

#### Protein digestion and MS/MS data collection

Protein samples were rapidly thawed and injected onto an integrated fluidics system containing a HDx-3 PAL liquid handling robot and climate-controlled chromatography system (LEAP Technologies), a Dionex Ultimate 3000 UHPLC system, as well as an Impact HD QTOF Mass spectrometer (Bruker). The protein was run over either one immobilized pepsin column at at 10°C (Trajan; ProDx protease column, 2.1 mm x 30 mm PDX.PP01-F32) at 200 mL/min for 3 minutes. The resulting peptides were collected and desalted on a C18 trap column (Acquity UPLC BEH C18 1.7mm column (2.1 x 5 mm); Waters 186003975). The trap was subsequently eluted in line with a C18 reverse-phase separation column (Acquity 1.7 mm particle, 100 x 1 mm^2^ C18 UPLC column, Waters 186002352), using a gradient of 5-36% B (Buffer A 0.1% formic acid; Buffer B 100% acetonitrile) over 16 minutes. Mass spectrometry experiments acquired over a mass range from 150 to 2200 m/z using an electrospray ionization source operated at a temperature of 200C and a spray voltage of 4.5 kV.

#### Peptide identification

Peptides were identified from the non-deuterated samples of FL and PX Sgk3 using data-dependent acquisition following tandem MS/MS experiments (0.5 s precursor scan from 150-2000 m/z; twelve 0.25 s fragment scans from 150-2000 m/z). MS/MS datasets were analysed using PEAKS7 (PEAKS), and peptide identification was carried out by using a false discovery based approach, with a threshold set to 1% using a database of purified proteins and known contaminants found in Sf9 cells (Dobbs *et al*, 2020). The search parameters were set with a precursor tolerance of 20 ppm, fragment mass error 0.02 Da, charge states from 1-8, leading to a selection criterion of peptides that had a −10logP score of 21.4.

#### Mass Analysis of Peptide Centroids and Measurement of Deuterium Incorporation

HD-Examiner Software (Sierra Analytics) was used to automatically calculate the level of deuterium incorporation into each peptide. All peptides were manually inspected for correct charge state, correct retention time, appropriate selection of isotopic distribution, etc. Deuteration levels were calculated using the centroid of the experimental isotope clusters. Results are presented as relative levels of deuterium incorporation and the only control for back exchange was the level of deuterium present in the buffer (81.5%). Differences in exchange in a peptide were considered significant if they met all three of the following criteria: : ≥7% change in exchange, ≥0.5 Da difference in exchange, and a p value <0.01 using a two tailed student t-test. All compared samples were set within the same experiment. To allow for visualization of differences across all peptides, we utilized number of deuteron difference (#D) plots. These plots show the total difference in deuterium incorporation over the entire H/D exchange time course, with each point indicating a single peptide. The mass spectrometry proteomics data have been deposited to the ProteomeXchange Consortium via the PRIDE partner repository (Perez-Riverol *et al*, 2019) with the dataset identifier PXD025327. The data analysis statistics for all HDX-MS experiments are in Supplementary Table 1 according to the guidelines of Masson et al (Masson *et al*, 2019).

