## Supplementary Figures for "*In vitro* reconstitution of Sgk3 activation by phosphatidylinositol-3-phosphate"

### Figure S1. Characterization of recombinant Sgk3.

**A**

*GluC digest (coverage = 48.4%; 55.7% coverage of kinase domain)*
*MS/MS of Sgk3 purified from Sf9 insect cells*

| Residue # | Location | ratio mod/base (fraction modified) |
|---|---|---|
| 21 | PX domain | ∞ (no base peptide detected) |
| 144 | PX-kinase linker | 0.42 (0.30) |
| 320 | activation loop | 0.27 (0.21) |
| 461 | turn motif | 0 (no modified peptide detected) |
| 486 | hydrophobic motif | 0 (no modified peptide detected) |

**B**

*GluC digest (coverage = 68.7%; 76.8% coverage of kinase domain)*
*MS/MS of Sgk3 purified from HEK293 cells*

| Residue # | Location | ratio mod/base (fraction modified) |
|---|---|---|
| 30 | PX domain | 0.02 (0.02) |
| 320 | activation loop | 0.06 (0.06) |
| 461 | turn motif | 0 (no modified peptide detected) |
| 486 | hydrophobic motif | 0 (no modified peptide detected) |

**C**

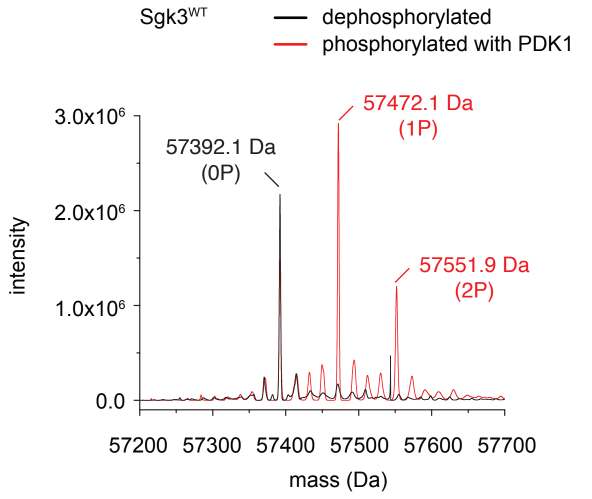


| Mass (Da) (expected) | Mass (Da) (observed) | Δmass (Da) | Matching modifications |
|---|---|---|---|
| 57391.8 | 57392.1 | +0.3 | none |
| | 57392.1 | +0.3 | none |
| | 57472.1 | +80.3 | 1P |
| | 57551.9 | +160.1 | 2P |

**D**

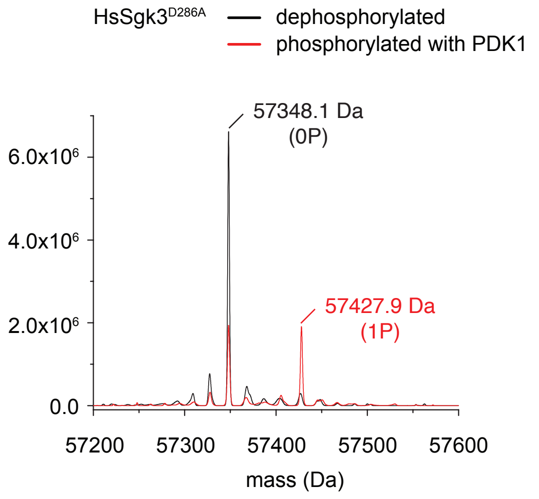


| Mass (Da) (expected) | Mass (Da) (observed) | Δmass (Da) | Matching modifications |
|---|---|---|---|
| 57347.8 | 57348.1 | +0.3 | none |
| | 57348.1 | +0.3 | none |
| | 57427.9 | +80.1 | 1P |

**E**

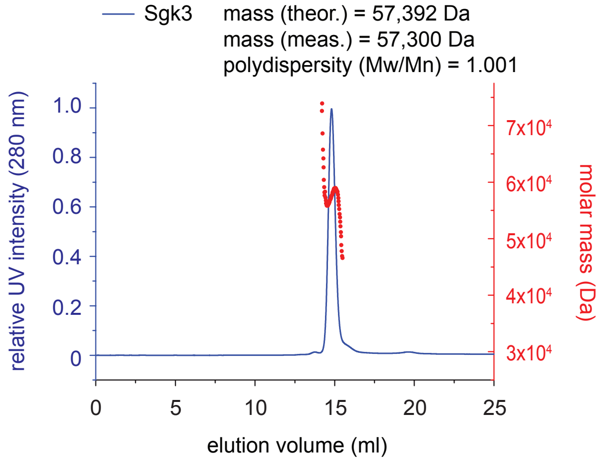


**F**

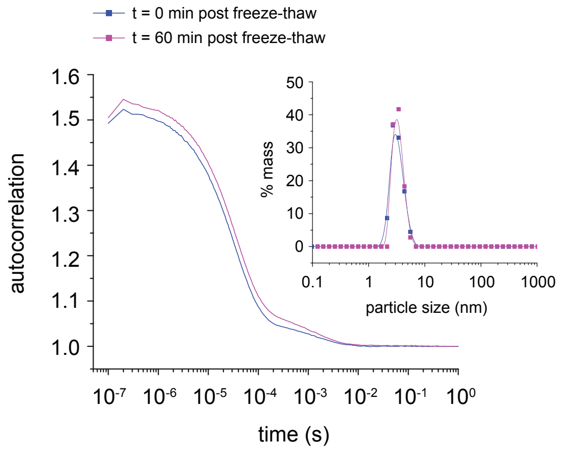


**G**

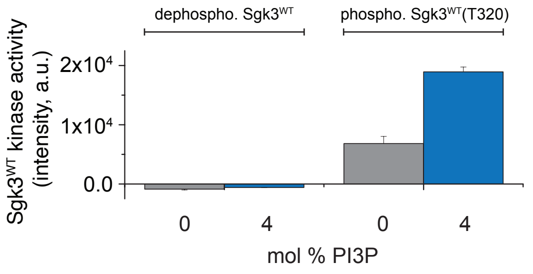

**Supplementary Table 1. Statistics for HDX datasets associated with this manuscript.**

| **Data set** | **FL Sgk3 0% PI3P** | **FL Sgk3 5% PI3P** | **PX Sgk3 0% PI3P** | **PX Sgk3 5% PI3P** |
|---|---|---|---|---|
| HDX reaction details | $\%D_2O$=81.5%<br>$pH_{(read)}$=7.5<br>Temp=18ºC | $\%D_2O$=81.5%<br>$pH_{(read)}$=7.5<br>Temp=18ºC | $\%D_2O$=81.5%<br>$pH_{(read)}$=7.5<br>Temp=18ºC | $\%D_2O$=81.5%<br>$pH_{(read)}$=7.5<br>Temp=18ºC |
| HDX time course (seconds) | 3, 30, 300, 3000 | 3, 30, 300, 3000 | 3, 30, 300, 3000 | 3, 30, 300, 3000 |
| HDX controls | N/A | N/A | N/A | N/A |
| Back-exchange | Corrected based on $\%D_2O$ | Corrected based on $\%D_2O$ | Corrected based on $\%D_2O$ | Corrected based on $\%D_2O$ |
| Number of peptides | 110 | 110 | 25 | 25 |
| Sequence coverage | 94.6% | 94.6% | 83.9% | 83.9% |
| Average peptide /redundancy | Length=13.9<br>Redundancy= 3.1 | Length=13.9<br>Redundancy= 3.1 | Length=12.2<br>Redundancy= 3.1 | Length=12.2<br>Redundancy= 3.1 |
| Replicates | 3 | 3 | 3 | 3 |
| Repeatability | Average StDev=1.9% | Average StDev=2.1% | Average StDev=1.6% | Average StDev=1.1% |
| Significant differences in HDX | >7% and >0.5 Da and unpaired t-test ≤0.01 | >7% and >0.5 Da and unpaired t-test ≤0.01 | >7% and >0.5 Da and unpaired t-test ≤0.01 | >7% and >0.5 Da and unpaired t-test ≤0.01 |

**Supplementary Figure Legends**

**Supplementary Figure S1**

**A)** GluC digest MS/MS phosphomapping of $Sgk3^{WT}$ purified from Sf9 insect cells.

**B)** GluC digest MS/MS phosphomapping of $Sgk3^{WT}$ purified from transiently transfected HEK293T cells.

**C)** Intact MS of dephosphorylated (black) and PDK1-phosphorylated (red) $Sgk3^{WT}$ protein.

**D)** Intact MS of dephosphorylated (black) and PDK1-phosphorylated (red) $Sgk3^{D286A}$ protein.

**E)** SEC-MALS profile of $Sgk3^{WT}$ indicating molecular weight and monodispersity.

**F)** DLS measurement of $Sgk3^{WT}$ in conditions of kinase assays measured after thawing (blue curve) and after 60 min incubation at RT (pink curve). Particle size distribution remains unchanged after 60 min incubation.

**G)** Kinase activity of dephosphorylated $Sgk3^{WT}$ vs. PDK1-phosphorylated $Sgk3^{WT}$ in presence of 100 µM Crosstide with 0% PI3P (grey bars) or 4% PI3P (blue bars). Error bars are the standard deviation of three independent experiments.

**Supplementary Table 1. Statistics for HDX datasets associated with this manuscript.**
